# High-resolution 3D imaging and topological mapping of the lymph node conduit system

**DOI:** 10.1101/752972

**Authors:** Inken D. Kelch, Gib Bogle, Gregory B. Sands, Anthony R. J. Phillips, Ian J. LeGrice, P. Rod Dunbar

## Abstract

The conduit network is a hallmark of lymph node microanatomy, but lack of suitable imaging technology has prevented comprehensive investigation of its topology. We employed an extended-volume imaging system to capture the conduit network of an entire murine lymph node (≈280,000 segments). The extensive 3D images provide a comprehensive overview of the regions supplied by conduits including perivascular sleeves, and distinctive “follicular reservoirs” within B cell follicles, surrounding follicular dendritic cells. A 3D topology map of conduits within the T cell zone showed homogeneous branching, but conduit density was significantly higher in the superficial T cell zone compared to the deep zone, where distances between segments are sufficient for T cells to lose contact with fibroblastic reticular cells. This topological mapping of the conduit anatomy can now aid modeling of its roles in lymph node function, as we demonstrate by simulating T cell motility in the different T cell zones.

## Introduction

Sophisticated immune responses are organized within the highly-structured microanatomy of lymph nodes (LNs) where stromal cell networks support the circulation, maintenance, and interaction of highly motile hematopoietic cell types on their continuous quest for cognate antigen (1-3). A key feature of the LN organization is the mesh-like network of fibroblastic reticular cells (FRCs) spanning the LN paracortex, the main homing zone for T cells (4, 5). FRCs organize LN microenvironments and control T cell life in many ways by providing survival signals, aiding migration, and restricting T cell activation (6, 7). They express the chemokines CCL19 and CCL21, important cues for motility, compartmentalization and retention of CCR7-expressing T cells, B cells, and dendritic cells (DCs) (3, 8, 9). In a similar fashion, FRCs appear to be involved in B cell homeostasis, by providing the B cell survival factor BAFF and contributing to CXCL13 expression (10, 11). LN expansion during immune stimulation is mediated by FRCs in synergy with DCs, which can trigger FRC stretching via interaction of CLEC-2 with podoplanin (12, 13). FRC destruction is part of the pathology of several devastating viral diseases, and directly affects the number and functionality of T cells (2, 7, 14). FRC networks also appear in tertiary lymphoid structures at sites of chronic inflammation underlining their central importance to immunobiology (15, 16).

Remarkably, FRCs construct a piping system that rapidly conducts incoming lymphatic fluid including tissue-derived antigens across the LN cortex (17-19). This conduit system consists of interconnected ‘micro vessels’ built of a central core of collagen fibers surrounded by a layer of microfibrils and a basement membrane enwrapped by FRCs, and channels molecules < 70 kDa from the subcapsular sinus (SCS) to inner LN compartments (17-20). In particular, inflammatory soluble mediators and cytokines can be shuttled directly to high endothelial venules (HEVs), specialized vessels for lymphocyte entry that are surrounded by perivascular “sleeves” formed by FRCs (5, 21-23). Intriguingly, the conduit network persists even if FRCs are temporarily lost, suggesting that it possesses structural integrity, while depending on FRCs for remodeling (10). Many questions remain concerning the heterogeneity of FRC populations, the exact mechanisms by which they regulate immunity, and the advantages of FRC-guided migration of T cells in a 3D space (7, 24). Our understanding of the structure of the conduit network remains limited due to the technical difficulty of capturing these delicate network structures within large tissue volumes (25). Previous approaches to studying the FRC network globally within LNs have relied on *in silico* computer models with pre-defined network properties (26-28), based on information from confocal images on a small scale (29, 30). Large-scale 3D imaging of entire networks has to date been hindered by the limitations of tissue penetration in standard microscopy, and restrictions in resolution of large-scale imaging techniques (31, 32). An additional complexity is that moving from small-scale measurements in 2D to large-scale measurements in 3D requires specialized non-trivial algorithms that often require custom computation by the operating lab to fit a particular purpose (32).

To provide a comprehensive picture of the LN conduit network we used a unique confocal block-face imaging system referred to as EVIS (<u>e</u>xtended-<u>v</u>olume imaging <u>s</u>ystem) (33, 34) and captured the conduit and blood vessel system of an entire murine LN. From the obtained seamless 3D images, we extracted a continuous topology map of the conduits in the T cell zone (TCZ) and quantified the network structure with the help of custom image processing tools. The obtained topology map permitted the assessment of 3D network parameters at unprecedented scale and served as a realistic template for *in silico* simulations of T cell motility. Our measurements revealed significant differences in conduit segment density between the deep and superficial TCZs, making it likely that T cells in the deep zone lose contract with the FRC network more frequently. We were surprised to find distinctive tracer accumulations in the B cell follicles, and we visualized the intriguing organization of the conduit-supplied spaces surrounding FDCs with new clarity. Our topology map provides a unique reality-based road map of the intricate 3D organization of the LN conduits that can be incorporated into the increasingly sophisticated theoretical models seeking to understand and predict complex immune processes within LNs (35).

## Results

### Extensive 3D imagery permits volume views of the continuous conduit network

Previously, studies of the LN conduit system have relied on microscopic images with limited depth information. By performing EVIS imaging at a voxel resolution of 1 µm we were able to capture a popliteal LN sized 850 × 750 x 900 µm in its entirety. Organ-wide anterograde labelling of the lymphatics and blood vessels was achieved by injecting wheat germ agglutinin (WGA) conjugated to different fluorophores into the footpad and the supplying blood vessel, respectively. The resulting 3D image permits detailed insights into the overall LN anatomy (Fig 1). As a 38kDa molecular tracer, WGA recapitulates the routes of lymph-borne molecules <70kDa through the LN. Strong WGA-labelling can be seen in the SCS and the medulla, thereby fully enclosing the LN. By virtually cropping the 3D volume (Fig 1 a), views of the interior organization (Fig 1, b) and the dense network of blood vessels running through the LN are revealed (Fig 1 c). The conduit network is most structured in the central TCZ (Fig 1 d), and is sparse in the B cell follicles, with only a few channels running beside any one follicle (Fig 1 e). The medulla is richly filled with WGA, providing a high staining intensity in the lymphatic sinuses, yet medullary cords, strands of parenchymal tissue that extend into the medullary space and are densely packed with cells (5, 36), are clearly distinguishable and contain at least one central blood vessel (Fig 1 f).

**Fig 1.**
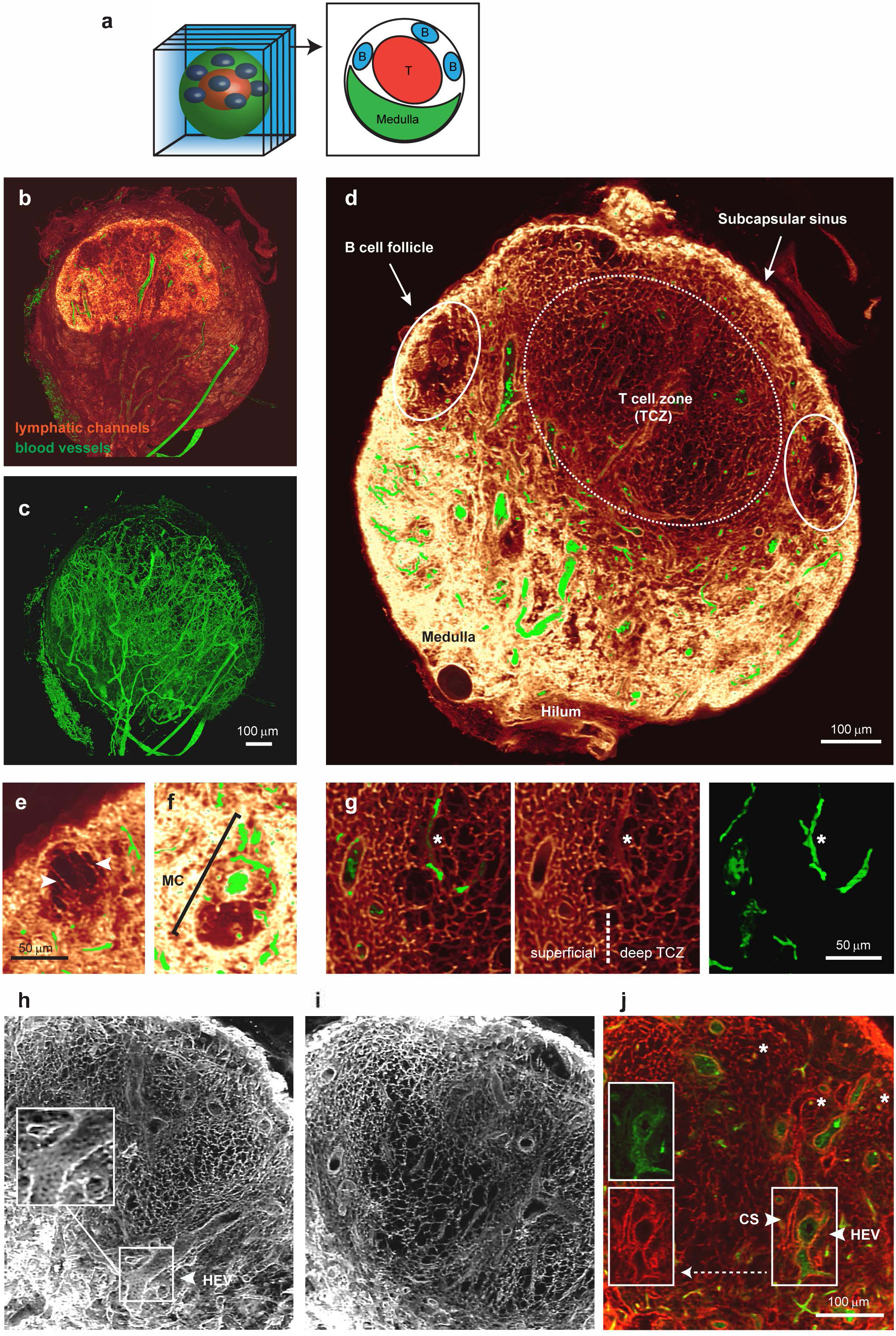
Detailed 3D images of the conduit network in a whole LN. EVIS imaging of an entire popliteal LN generated a 3D volume image of which interior slices can be viewed individually (**a**). 3D image reconstruction of the entire LN shows (**b)** lymphatic channels filled with the tracer molecule WGA (red glow) together with dextran-labelled blood vessels (green), or the blood vasculature alone (**c**). An interior view of 20 µm thick optical sections (**d - i**) and a 1 µm slice (**j**) permits detailed insights into the LN architecture. A cross-section of the LN displays the location of cell-specific zones (**d**), while close-ups reveal anatomical details including the arrangement of long conduits descending from the SCS at the edges of a B cell follicle (arrowheads, **e**), a medullary cord (MC) with a central blood vessel situated amongst the WGA-filled sinuses of the medulla (**f**), and the transition from dense to sparse conduit networks in the superficial to the deep TCZ (**g**). The conduit network forms a highly organized grid within the TCZ (white, **h, i**; red, **j**) interspersed with cortical sinuses (CS, arrowhead, **j**) and blood vessels (green) including HEVs, which are closely surrounded by cells displaying a cobblestone-like morphology (arrowhead, **h, j**). Besides larger blood vessels, small blood vessels are frequently enclosed by conduits (asterisks, **g, j**). Image rendering was performed in Voxx (**a – i**) and ImageJ (**j**). See also S1 Fig, S1 Video, and S2 Video.

Within the TCZ the conduit network appears most dense in the superficial and interfollicular zones, while a sparser network structure becomes apparent within its center (Fig 1 d, g-j), consistent with previous definitions distinguishing the deep TCZ from surrounding regions (37). Particularly strong staining could also be observed around HEVs and smaller blood vessels, which both appear surrounded by a sleeve contiguous with the conduit network (Fig 1 g-j). However, intraluminal staining of blood vessels with lymph-derived WGA was not observed (Fig 1 g-j, S2 Fig). Cortical sinuses (38) also display strong labelling, but can be distinguished from blood vessels by lack of blood vessel-specific WGA-staining (Fig 1 j) and their continuity with the medullary sinuses, a feature that becomes evident in animations of the 3D dataset (S1 and S2 Video). Interestingly, while it was previously reported that conduits are primarily focused on HEVs, we observed in our 3D images that conduits frequently terminate on cortical sinuses, which are often located in close proximity to blood vessels (S1 Fig). Examining tissue sections using conventional immunofluorescence microscopy confirmed that conduits connect to cortical sinuses made up of LYVE-1+ lymphatic endothelial cells (S1 Fig). Together, these images demonstrate that the conduit system connects the SCS with cortical sinuses that drain into the medulla, as well as the perivascular sleeves surrounding blood vessels including HEVs, thereby providing a continuous piping system for incoming lymphatic fluid (S1 and S2 Video).

### Quantification of the conduit network topology

The availability of an extensive 3D volume image of the continuous LN conduit network permits quantification of its network statistics at unprecedented scale and provides an exciting opportunity for the realistic modeling of T cell motility. We previously imaged and quantified the blood vessel system of a mesenteric LN using a set of custom-developed image processing and analysis tools (33), and now applied these tools to perform large-scale 3D analysis on the conduit network. The image processing consists of a number of steps including thresholding and skeletonization, which transform the pixel-based image data into a 3D topology map. The topology map describes the network as a system of connected tubes and enables a direct read-out of network parameters (Fig 2). In order to study the network topology of the conduit system in the central paracortical TCZ and its implications for T cell biology, the extraction procedure was optimized to best capture the network in this region (Fig 2 a-c). A limitation of this process was posed by the occurrence of continuous tracer-labeled spaces fully enclosing large blood vessels, such as HEVs (S2 Fig, S3 Video), identified previously as perivascular sleeves (5).

**Fig 2.**
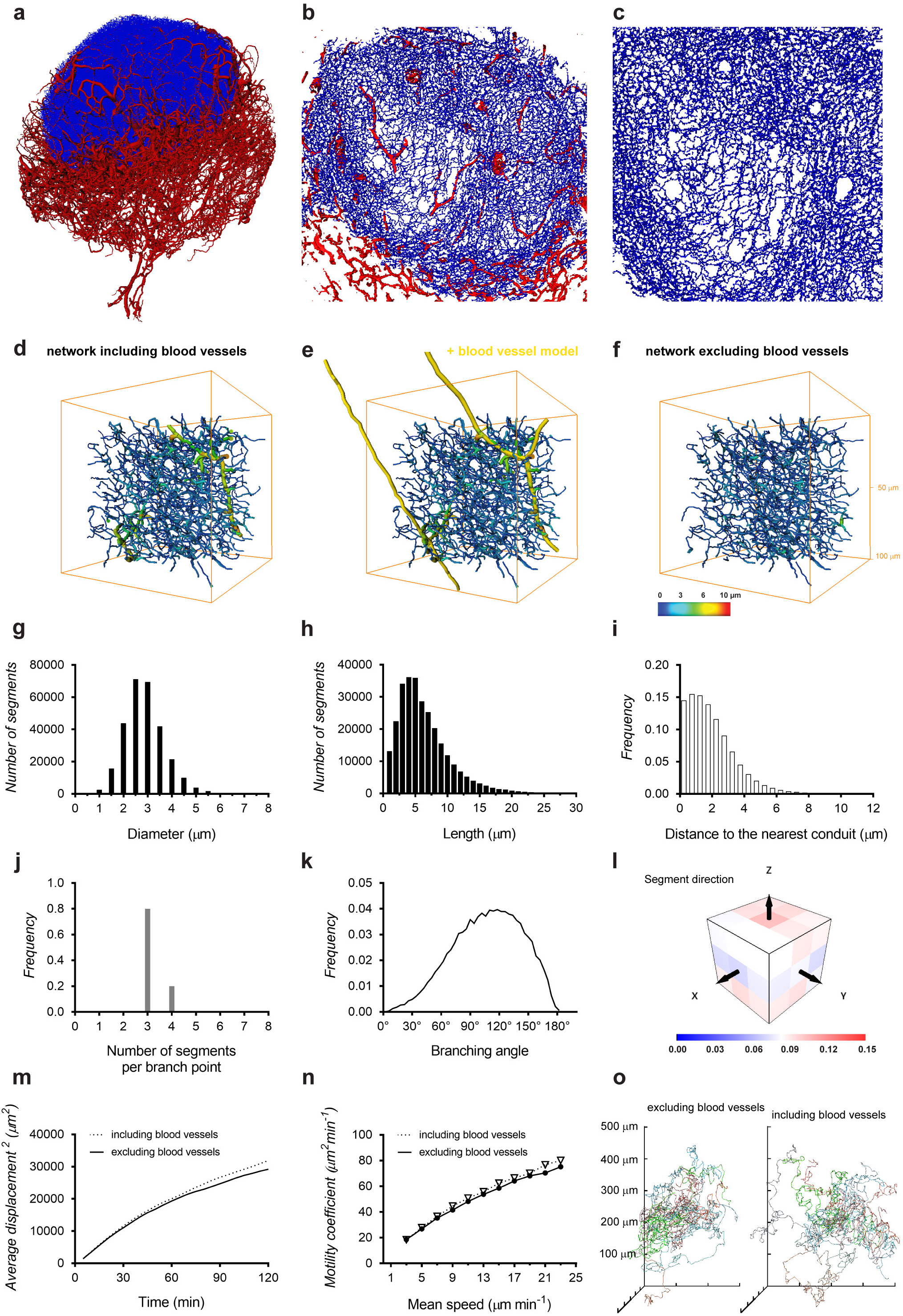
Network topology of the LN conduits in the TCZ. Deploying custom-developed image-processing tools, a description of the conduit network in terms of nodes and links was generated from the 3D image data and used to estimate network parameters. 3D projections of the blood vasculature (red) and the conduit network in the TCZ (blue) as a whole (**a**) and magnified views of the TCZ (**b, c**) expose the complexity and the high level of detail in this dataset. The full conduit data includes large segments with diameters over 5 µm (**d**), but these represent blood vessels as the overlay with the blood vessel model (yellow) indicates (**e**). A blood vessel-free conduit network (**f**) was obtained by removing the majority of blood vessels from the 3D image prior to the network extraction in a semi-automated process. This TCZ conduit network excluding blood vessels was employed to calculate the distribution of segment diameters (**g**), lengths (**h**), the branching pattern (**j**), branching angles (**k**), and segment orientation (**l**), while the full dataset including blood vessels was used to calculate the minimum distance to the nearest conduit (**i**). Simulation of T cell motility utilizing these conduit data provides the cell displacement at a mean speed of 13 µm min-1 (**m**), motility coefficients for different speeds (**n**), and a spider-plot representation of migration paths in a network with and without blood vessels (**o**). See also S2 Fig, S3 Fig and S3 Video.

This feature of the conduit network provided an obstacle for the skeletonization process (S3 Fig) and required us to adapt our image processing strategy. We overcame this problem by utilizing the co-stained blood vessels and subtracting the segmented blood vessel image data from the conduit image. In our previous study (33), we found blood vessels in the LN typically have diameters between 4 and 87 µm, while diameters of conduits are reported to lie in the range of 1 to 2 µm (17, 19, 20, 39). By removing the blood vasculature from the conduit data, vessels of the size of blood vessels could be effectively excluded (Fig 2 d-f). The resulting ‘clean’ conduit network contained 282,716 segments with a mean diameter of 2.9 µm and an average length of 6.5 µm (Fig 2 g, h; Fig 3 h, i). Within the TCZ conduit network, spanning a volume of about 0.079 mm^3^, the conduit segments had a combined length of 1.84 m and a density of 3.54·10^6^ segments mm^-3^ (Fig 3 f, j). To obtain a measure of spacing in the network, we applied an algorithm that measures the distance to the nearest conduit segment starting from a regular fine grid of points located in the LN volume (33). This calculation revealed that the majority of locations in the LN TCZ lie within a very short distance of the nearest conduit (< 4 µm, 90.9%) (Fig 2 i). Overall, the conduit network displayed an even branching pattern, with the majority of branching points representing bifurcations and branching angles centered around 120° (Fig 2 j, k). The segment orientation had no observable bias in direction (Fig 2 l).

**Fig 3.**
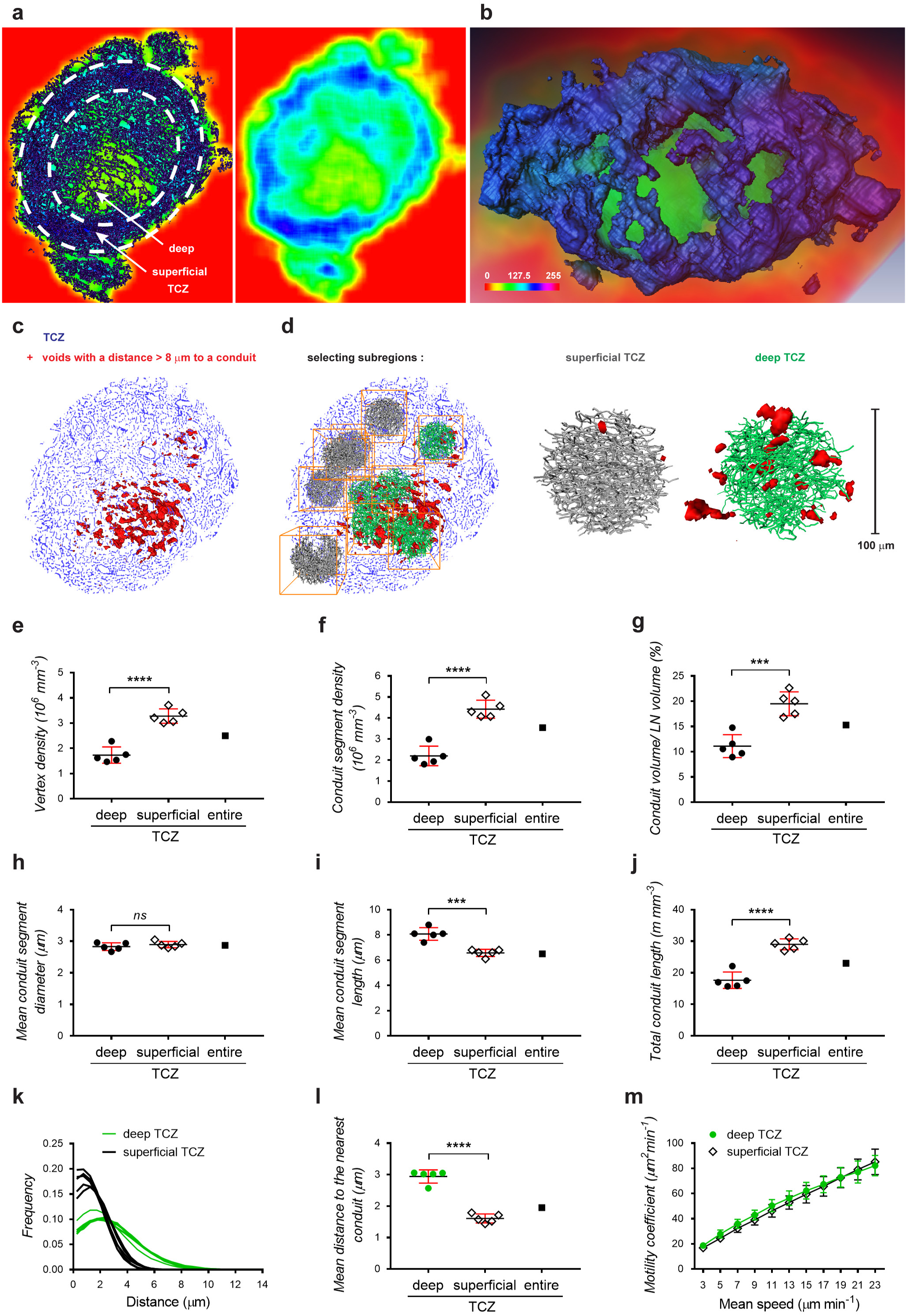
Comparison of conduit network parameters in the deep and superficial TCZ. Differences in conduit density between the deep and superficial TCZ can be visualized by averaging and color-coding pixel densities over small image volumes in a ‘moving average’ display, shown as a rainbow spectrum (**a, b**). A cross-section of the moving average display exposes how dense regions in the periphery of the LN (blue) surround an inner region of lower conduit density (green), directly representing dense or sparse occurrence of conduit segments in the corresponding section of the conduit network image (dark blue, left panel), respectively (**a**). Volume rendering (**b**) of the entire TCZ using this approach shows the dense superficial zone (blue) enclosing a central region of sparse conduits (green). Alternatively, the TCZ conduit map was employed for calculating the distances to the nearest conduit and voids with a distance of over 8 µm were displayed in red, indicating larger distances within the deep TCZ as opposed to outer regions (**c**). From these two zones 10 subregions were selected for comparative analysis (**d**); including the number of vertices (**e**), the number of conduit segments (**f**), the conduit volume (**g**), conduit segment diameters (**h**), conduit segment lengths (**i**), and the combined conduit length (**j**). The distributions of the minimum distances to the nearest conduit (**k**) and the average minimum distance (**l**) further exemplify the larger spacing within the deep TCZ. Simulation of T cell motility predicts similar motility coefficients within the deep TCZ and the surrounding superficial zone (**m**). Data are from one experiment (each point represents one 100 µm subregion, N = 10) and plots show means ± SD. **** p < 0.0001, *** p < 0.001, ns = not significant, Student’s t-test. See also S4 Fig.

We then tested how this network topology would predict the migration of T cells in 3D, when stimulated T cells are restricted to migrating along the network segments, as if in continuous contact with FRCs. We simulated the paths of a large number of cells on the extracted conduit network to calculate the coefficient of motility (*C*_*m*_) as an index of dispersal rate in 3D space, representing the rate at which T cells can scan a volume of paracortex for the presence of cognate antigen. In these simulations, we used values of mean speed in the range typically measured by intra-vital microscopy (40-44). The average displacement of cells (Fig 2 m) at T=60 min was used to calculate *C*_*m*,_ (Fig 2 n), and the correlation we generated between mean speed and *C*_*m*_ was broadly consistent with values previously measured *in vivo* (41). The average displacement and the corresponding *C*_*m*_ values are not significantly increased when blood vessels are included in the analysis (Fig 2 m, n), but cell tracks show a slight variation due to the availability of the blood vessels and the sheaths that often surround them as additional migration paths (Fig 2 o).

### Topology differences in the deep and superficial TCZ

It was evident in the 3D conduit image and the topology map that the conduit network in the TCZ is not homogeneous, but displays different densities in the superficial and deep zone (Fig 3), concordant with previous descriptions (37). After coloring regions based on their segment density, it is possible to visually distinguish the deep TCZ, containing a rather open mesh, from the superficial zone, which is characterized by a dense network of conduits and fully encloses the spherical central T cell region (Fig 3 a, b). In a different approach to visualizing the variable spacing in the network, distances to the nearest conduit segment were measured in 3D and locations further than 8 µm from any conduit were displayed in red (Fig 3 c, d). An accumulation of red voids is located centrally in the deep TCZ, while they were absent from superficial locations. To quantify these regional differences, 5 spherical subregions with a diameter of 100 µm were selected from the superficial and deep TCZ each and examined using the topology toolset (Fig 3 d). The deep TCZ contained significantly fewer vertices, segments, and a smaller conduit volume per region than the superficial zone, confirming visually observable differences in conduit density (Fig 3 e-g). While the conduit diameters in both locations showed no measurable difference, individual segment lengths were considerably shorter in the superficial zone, yet the combined conduit length of all segments was longer than in the deep TCZ (Fig 3 h-j). In summary, the deep TCZ can be perceived as a stretched version of the conduit network in superficial areas. As a result, cells in the deep TCZ have a 50% greater mean distance to the nearest conduit segment (Fig 3 l), reaching distances well beyond the cell diameter of a murine lymphocyte (2.5-3 µm) (45), and making it unlikely that cells in this region are in contact with a conduit segment at all times. In contrast, distances measured in the superficial zone would allow nearly continuous contact with the network (conduit distance < 4 µm: 73.4% in the deep TCZ vs. 96.8% in the superficial TCZ; conduit distance < 6 µm: 92.2% deep vs. 99.9% superficial) (Fig 3 k).

We then used our simulations of T cell migration to predict motility coefficients separately in the superficial and deep zones, assuming that T cells remained in contact with the conduit network. The calculated 3D motility coefficients gradually increased with the speed of migration but there were no significant differences in motility coefficients between the zones (Fig 3 m).

With respect to potential specialised immune functions within the different TCZs, we noted differences in the distribution of proliferating T cells in 2D sections of resting LNs. Ki-67+ proliferating cells were often found in close proximity to the conduit network, and seemed more frequent in the peripheral TCZ than the deep T cells zone (S4 Fig), reinforcing the possibility that close cell contact (or the cues they provide) is important for T cells in the superficial TCZ.

### Conduit organization in B cell follicles

EVIS imaging of WGA-perfused LNs led to the unexpected observation of distinctive tracer accumulations inside B cell follicles. Compared to the dense conduit network in the TCZs, conduits are very rare in the B cell zones (Fig 4 a, b), although a small number of conduits could often be visualized descending directly from the SCS, consistent with channels previously referred to as follicular conduits (39). However, unexpectedly we also observed distinctive WGA tracer accumulations within the B cell regions, appearing as discrete multilobular spaces reminiscent of ‘honeycombs’ that are connected to the SCS and each other via follicular conduits (Fig 4 a, b, S4 Video), occasionally aggregating into larger contiguous cavities. Hence these clusters appear as striking dense accumulations of WGA tracer within B cell zones that are otherwise relatively devoid of conduits (Fig 4 g, h). To test how these WGA-filled spaces relate to the location of follicular dendritic cells (FDCs), we used multicolor immunohistochemistry to identify FDCs in WGA-perfused LNs. The FDC marker CD21/CD35 co-localized with the observed deposits of WGA tracer (Fig 4 c), confirming that the spaces we visualized surrounded and intercalated with FDCs deep within B cell follicles. Additional stains using collagen I to visualize conduit channels confirmed the transport of WGA through follicular conduits and deposition of WGA on FDCs (Fig 4 d). Moreover, the arrangement and morphology of FDCs within the B cell follicle, as shown by co-staining with collagen I and a B cell marker, closely mirrors the location of the spaces typically filled by WGA (Fig 4 e, f). High-resolution confocal image stacks revealed some diversity in the spaces where the WGA tracer accumulated within the follicles (Fig 4 g, h, S5 Video). As well as the almost spherical structures of ∼30 µm diameter that were brightly labeled, we noted weaker WGA tracer accumulation in adjacent honeycombed regions (Fig 4 h, arrows), consistent with the various shapes of FDC clusters (Fig 4 e, f). We also noted that the WGA signal inside B cell follicles was not as abundant in the 2D frozen sections (Fig 4 c, d) compared with our 3D data (Fig 4 a, b, g, h), suggesting that tracer may be washed off during frozen section preparation while being retained in the PFA-fixed and resin-embedded LNs we used for 3D imaging.

**Fig 4.**
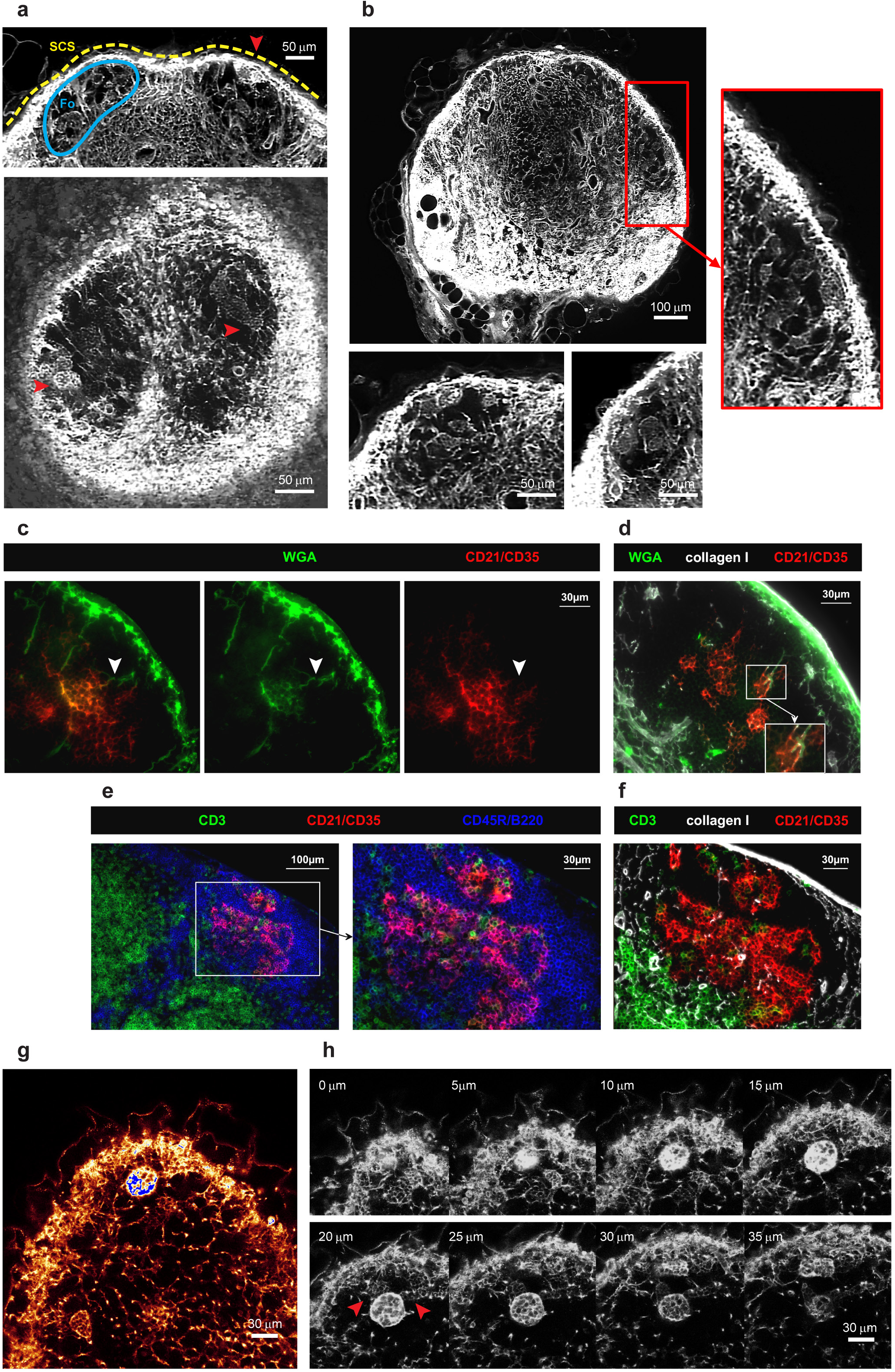
The follicular conduit. 3D EVIS images of a popliteal LN with WGA-labelled conduit paths contain brightly labelled multilobular spaces (red arrowheads) within the otherwise unstained B cell follicles (Fo) underneath the SCS that can be repeatedly seen in 3D projections of 20 µm thickness (**a**) and 2D image slices (**b**). In immuno-labelled tissue sections of WGA-perfused inguinal LNs, WGA (green) is found in follicular conduits descending from the SCS (white arrowheads) connecting to cellular clusters expressing the FDC-marker CD21/CD35 (**c, d**). The morphology and location of WGA-labelled cell clusters within B cell follicles (**d**) are generally consistent with the anatomy of FDCs in these regions (**e, f**) as co-staining with markers for B cells (CD45R/B220), T cells (CD3), and collagen I confirms. High-resolution confocal images (with a voxel resolution of 0.36 × 0.36 × 1 µm) of a WGA-perfused popliteal LN provide insights into the staining pattern within and around the WGA accumulations (**g**), and show a particularly bright cluster in several z steps (**h**) directly neighboring spaces with weaker labelling (red arrowheads). Images are representative of at least 6 LNs (from N = 5 mice) in which multilobular cell clusters could be observed. See also S4 Video and S5 Video.

## Discussion

We set out to map the conduit network across an entire LN to enable measurements of its topology. Here we present extensive 3D imagery of the conduit channel system of a whole LN, permitting detailed insights into the conduit organization and its connectivity with blood and lymphatic vessels. We also provide a continuous reality-based computer representation of the TCZ conduits to enable large-scale quantification and downstream use in computer models of immune processes.

The conduit system has an intricate spatial relationship with the blood vessels and the lymphatic sinuses. Besides stabilizing the organ structure through their scaffold-like organization, conduits are thought to provide a short cut between incoming lymph and HEVs (18, 19). Although there is evidence that some molecules such as chemokines can gain access to the HEV lumen through the conduit network (18, 19, 22), this may depend on transport through endothelial cells by transcytosis (46). We did not observe significant intra-vascular staining with WGA supplied into the conduits, implying that this 38 kDa molecule did not readily have access to the vascular lumen but was instead retained in a perivascular sleeve. Our 3D imagery therefore supports the concept that conduit segments descend from the SCS, branch through the paracortex, and richly supply perivascular sleeves including those surrounding HEVs. Notably, these perivascular sleeves represent the first space encountered by cells exiting the bloodstream, and suggest an under-explored role for the conduits, in conveying molecules directly from the SCS to lymphocytes and antigen-presenting cells that have just entered the LN from the blood. These data also lead us to conclude that these regions do not represent major sites of lymph drainage into the vasculature. Instead, we observed conduits frequently terminating in lymphatic sinuses that are blind-ended invaginations of the medullary sinuses, which is likely to provide the necessary outlet for accumulating lymph and potentially aids cell egress at these locations (47).

Analysis of over 280,000 conduit segments in the 3D topological model we generated revealed a homogeneous branching pattern, with bifurcations being the most prevalent branching structure and no more than 7 segments meeting at any one point. The level of connectivity that we measured between neighboring nodes is slightly lower than that measured by Novkovic, Onder (30), who note the presence of highly connected nodes with more than 12 edges, based on confocal images of CCL19-expressing cells across a small fraction of the paracortex. The data presented here represent network parameters from an entire TCZ providing a clear advantage to previous extrapolations from small LN regions. In contrast to their cell-based graph network, our conduit model provides a road map of the collagen-bearing conduit channels that the FRCs ensheath, including their exact lengths and orientation. It is possible that two or more conduit branch points in our model fall within the area of one cell body, to account for some of the differences in network topology, yet our data do not support the prevalence of highly connected nodes as they report. New opportunities are now likely to arise by combining the techniques we developed with those established for imaging FRC cell bodies, for example to track the structure of entire conduit networks in response to immune stimuli or disturbance of FRC network integrity, phenomena that have only recently begun to be explored (10, 13, 14, 30).

A striking feature that is visually obvious in our 3D images is the variation in conduit network density between the deep and superficial T cell regions. The topology map of the TCZ conduits we generated allowed us to quantify significant differences in conduit segment density, segment length, and inter-segment gap size between both zones. Our 3D imaging data are therefore consistent with several studies that previously identified a structural inhomogeneity within the TCZ in the LN paracortex. While the deep zone has been described as loosely interspersed with a network of FRCs and conduits, the peripheral zone was noted to contain a much denser mesh and a higher abundance of HEVs (37). The superficial TCZ (48) has also been referred to as the cortical ridge (37, 49), or the peripheral T cell region (50), and is continuous with the interfollicular regions between the B cell follicles closer to the SCS (51, 52). Although the biological significance of this structural segregation is still unclear, independent reports have pointed to an asymmetry in cell positioning in both zones. Naïve T cells tend to occupy the deep TCZ, whereas memory T cells preferentially locate to the superficial zones, and innate effector cells can often be found in the interfollicular regions (37, 53, 54). Similarly, subtypes of resident and migratory DCs seem to preferentially locate to either the deep, the superficial, the interfollicular zones, or regions close to the medulla (49, 50, 55). It has also been frequently observed that following immune challenge T cells cluster in peripheral regions or locations close to the medulla (50, 51, 56-58). It is therefore intriguing to note that Ki67-expressing cells in the resting LNs we examined were often located in very close proximity to a conduit, and tended to localize to the periphery of the TCZs (S4 Fig). IL7-production may be higher in the peripheral TCZ (59), and close proximity to the FRC network might increase access to homeostatic survival and growth factors for memory or recently-primed T cells.

Our measurements of conduit density in the deep and superficial TCZs led us to conclude that while T cells within the superficial zone could remain in almost continuous contact with FRCs wrapped around the conduits, the larger gap size in the deep T cells zone does not guarantee simultaneous contact for all T cells in this region. While *in vivo* imaging studies suggested that T cell motility is generally bound to the FRC network (43, 60), T cells were observed to occasionally leave the FRC paths and migrate perpendicular to the FRC scaffold. Recent data established that T cells migrate in a sliding manner on the FRC network and suggest that fast scanning rates are achieved through low adhesiveness to the FRC substrate (43, 61). Our 3D data confirm that a dense continuous network is present to support the migration of cells across the TCZ, but the more open topology in the deep TCZ substantially increases the likelihood of an occasional loss of contact. Interestingly, theoretical studies of the FRC network have concluded that the odds of a cognate T/DC encounter are in fact not significantly increased by confining migration to a network (25-28).

When we used our 3D topology map of the conduit network as pathways to simulate FRC-bound T cell migration, we observed an increase in the coefficient of motility as velocities increased across the range commonly measured *in vivo* (40, 41), confirming that higher velocities translate to faster scanning rates. However in these simulations, where T cell migration was solely restricted to the paths represented by the conduit network, we could not detect substantial differences in the coefficients of motility between the deep and superficial TCZs at any particular velocity. This may relate to the fact that although the density of the conduit networks differs in these two zones, their branching topology is very similar, with the network in the deep zone effectively representing a “stretched” version of that in the superficial zone. Some previous *in vivo* measurements recorded higher T cell velocities in the deep TCZ compared to peripheral zones, which implies that motility coefficients could differ accordingly in these regions *in vivo* (44, 62). Future models of T cell migration will now be able to incorporate our measurements of conduit network topology to model the conduit/FRC-guided component of T cell motility, as we have shown here. However, accurate models will benefit from incorporating additional factors, including the effect of chemokinesis and chemotaxis (63) especially driven by CCL19 and CCL21 (3, 57); the need for T cells to migrate around obstacles (26) including each other (64); and binding to DCs (44), as well as external factors such as confinement affecting the mode of cell migration (43, 61).

In summary, our data provide quantitative support for the concept that the FRC and conduit network in the paracortex are arranged in a way that supports different processes in spatially distinct functional zones.

In the B cell follicles, conduits have previously been found similar in diameter and particle size exclusion (molecules >70 kDa) to those in the TCZs, although they don’t span the B cell follicle but descend as sparse short parallel channels from the SCS to converge with FDCs in the center of the follicle (39). Here we confirm this topology of the follicular conduits, but we also show that they supply a space surrounding the FDCs, that we propose be termed “follicular reservoirs”. These honeycombed spaces surrounding FDCs are remarkably well defined and can easily be distinguished in 3D images from the voids of unstained cells surrounding them in the follicles. While these structures could be easily identified in all PFA-fixed and LR white-embedded preparations of whole LNs, staining was frequently lost in acetone-fixed cryosections, suggesting that the fluorescent tracer is in soluble, unbound form within the follicular reservoirs. This phenomenon may also explain why these structures have not been seen in this clarity in previous studies. Our methods to label and preserve the material within the follicular reservoirs open the way for future studies to identify all the cell populations involved in their formation, to clarify the mechanisms by which molecules from the SCS accumulate within them, and to track the changes they undergo during immune activation and germinal center formation.

The identification of follicular reservoirs supplied directly by fluid from the SCS is important when considering the supply of antigen to B cells (39, 65). While free diffusion limits the speed at which soluble antigen can reach the deeper follicular region from the SCS, the follicular conduit allows these materials to be rapidly channeled directly to B cells and FDCs in the center of the follicle, which have been shown to readily take up small non-complexed molecules (66, 67). In the presence of local Ig or small complement molecules soluble antigen could then be complexed and retained by FDCs to fulfill the prerequisite for sustained B cell activation (66, 68). We suggest that the follicular reservoirs we have identified are likely to play a pivotal role in this process. In addition, the follicular conduits may enable FDCs to access signaling molecules < 70kDa delivered from incoming lymphatic fluid or cells near the SCS, in order to respond rapidly and directly to external stimuli.

It is important to note that the precise roles of the conduit system in distributing molecules to different LN compartments remain unclear. Several groups reported that DCs and B cells can obtain antigen directly from the conduits, providing a fast antigen delivery system that extends deeply within the LN (19, 20, 67, 69). However, Gerner, Casey (50) have challenged this prevailing view, concluding that antigen dispersal to DCs and subsequent T cell stimulation is dominated by conduit-independent diffusion. Instead the conduit system may simply enable equilibration of fluid, with a subsidiary role in transporting signaling molecules (50). Thierry, Kuka (70) recently provided additional support for the conduit system acting as a drainage system that allows IgM produced in the parenchyma to readily exit the LN and assure a rapid response to infection. Specifically localizing these different immune processes with respect to the spatial variations in the conduit network will help improve our ability to control and manipulate immune responses (50, 71).

In summary, the data reported here present the first reality-based description of the conduit network across an entire murine LN paracortex. The extracted topology network provides a useful substrate for theoretical models of LN biology (35), such as 3D motility models and models of fluid distribution (29), as well as providing new insight into the structure of a network that is crucial to many immune functions.

## Materials and Methods

### Mice

All animal work was performed in accordance with the guidelines and the requirements of the New Zealand Animal Welfare Act (1999) and approved by the University of Auckland’s Animal Ethics Committee. C57BL/6J mice were purchased from The Jackson Laboratory. Experimental protocols employ 9-22 weeks old male C57BL/6J mice housed in the conventional animal facility unit at the School of Biological Sciences at the University of Auckland under environmentally controlled conditions (temperature and humidity) and a 12:12-h light/dark cycle. Animals were group-caged in transparent IVC cages with wood-chip bedding and environmental enrichment, in close proximity to other cages so that auditory, visual and olfactory stimulation was present. We assessed animals daily for health and welfare, and access to food and water.

### Tissue preparation for EVIS imaging

For *in vivo* staining of murine LNs we used Alexa Fluor 488, 555, TMR or 647 conjugated WGA (wheat germ agglutinin, Invitrogen), TMR-conjugated 2000 kDa dextran (Invitrogen) and anti-LYVE-1 antibody (R&D Systems) that was fluorescently conjugated using the Alexa Fluor 488 Antibody Labelling Kit (Invitrogen). For the labelling of LN conduit paths fluorescently conjugated WGA was used as an anterograde tracer. After brief anesthesia, 50 µl of WGA-Alexa Fluor 488 (1mg/ml) were injected into the footpad of C57BL/6 mice and let to circulate for 30-60 minutes. This was followed by labelling the blood vascular system using a sequence of 1 ml fluorescent WGA-TMR (50 µg/ml at 20 µl/min) and 1 ml 2000 kDa dextran-TMR (500 µg/ml, Invitrogen)/2.5% gelatin mix (at 50 µl/min) in a post mortem local perfusion technique as described earlier (33). Excised popliteal LNs were fixed in 4% PFA, 3% sucrose at 4°C overnight before embedding in stable resin for EVIS imaging. Resin embedding was carried out by first dehydrating the tissue and infiltrating with LR white (hard grade, ProSciTech) followed by curing for 6 hours at 60°C as previously described (33). The observable tissue shrinkage that occurs during this process was estimated to be 20%.

### Tissue staining and conventional confocal microscopy

To achieve triple staining of the blood vasculature, conduit channels, and lymphatic vessels in popliteal and inguinal LNs, anaesthetized C57BL/6 mice were first injected with 50 µl anti-LYVE1-Alexa Fluor 488 antibody (20 µg/ml, R&D Systems) into the rear right hock, an alternative injection site to the footpad which is less invasive while allowing strong labelling (72). After 8 hours, 50 µl WGA-Alexa Fluor 647 (1 mg/ml, Invitrogen) were injected in the same site and after a circulation period of 1 hour, the blood system of the whole body was labelled by injecting 100 µl of WGA-Alexa Fluor 555 (5 mg/ml, Invitrogen) with 10 µl Heparin (100 units/ml) into the tail vein or vena cava of the anaesthetized mouse for a duration 2 minutes. Freshly excised murine tissue was fixed in 4% PFA, 3% sucrose at 4°C overnight and embedded in LR white resin (medium grade, ProSciTech) for confocal imaging as described above. Standard confocal microscopy was performed using a Leica TCS SP2 equipped with a Leica HCX APO L 40.0×0.80 W UV water objective (Leica microsystems) at a voxel resolution of 0.36 × 0.36 × 1 µm.

### Immunohistochemistry

Freshly excised popliteal or inguinal LNs were snap frozen in O.C.T. compound (Sakura Finetek) and sectioned into 7 µm thick tissue sections. A protocol for multicolor immunohistochemistry established by Lloyd et al. (73) was adopted for immunostaining using up to four labels. These include antibodies against LYVE-1 (R&D Systems, clone 223322), collagen I (Abcam), laminin (Abcam), CD21/CD35-Biotin (Biolegend, clone 7E9), Ki-67 (Biolegend, clone 16A8), CD3e (BD Pharmingen, clone 500A2), and CD45R/B220 (BD Pharmingen, clone RA3-6B2). Primary antibodies were detected with Alexa Fluor 488, 555, or 647 conjugated goat secondary antibodies or Streptavidin (Invitrogen) and nuclei labelled with DAPI (Invitrogen). Stained immunohistochemistry sections were mounted using ProLong Gold Antifade reagent (Invitrogen) and photographed on a Nikon Eclipse Ni-U epifluorescence microscope (Nikon Instruments) using a SPOT Pursuit 1.4MP monochrome camera (Scitech). Acquired images were pseudo-colored, processed, and superimposed employing the Cytosketch software (Cytocode Limited).

### EVIS imaging and image processing

Extended-volume confocal imaging (EVIS) is a confocal block-face imaging method that can capture large 3D regions of fluorescently labelled tissue up to several millimeters thick at a pixel resolution of up to 0.5 µm. In an iterative process, a resin-embedded sample is moved between a confocal laser scanning microscope (TCS 4D CLSM, Leica) and a precision miller (Leica SP2600 ultramill, Leica) both mounted on a high-precision three-axis translation stage (Aerotech, US) and controlled by imaging software written in LabVIEW™ (National Instruments), whereby previously imaged sections are removed in between imaging rounds as previously described (33). Image acquisition was performed using an Omnichrome krypton/argon laser (Melles Griot) for sample illumination, a 20x water immersion lens (HC PL APO, 0.70 NA, Leica), 4x line averaging, and an image overlap of 50%. Individual 8-bit (grayscale) images acquired at ‘1 µm pixel resolution’ contained 512 × 512 pixels covering an area of 500 × 500 µm, providing a pixel resolution of 0.98 µm. By acquiring successive images at a z-spacing of 1 µm, an isotropic voxel size of (1 µm)^3^ was achieved. Precise xyz-registration of the acquired image stack in conjunction with custom-designed image processing and assembly software (LabVIEW™, (33, 34)) enables the composition of seamless 3D images. As part of this process, individual images underwent background correction, deconvolution, and denoising, before being merged into x-y mosaics and assembled into a 3D volume image. To further improve the quality of the generated 3D images and reduce the fluctuation of signal intensities between individual z planes across the 3D image stack, we employed an equalization algorithm to adjust the average image intensity in z direction. Rather than having a fixed target for correction, a variable (‘smoothed’) ideal intensity was used for each z plane, to account for the changing diameter across the spherical LN sample. Using the formula below, a correction factor *f*(z) was obtained for each z plane and multiplied with the pixel intensities on the respective plane to create an equalized image. The corrected 3D image displayed a significantly reduced intensity variation and was better suited for image analysis.

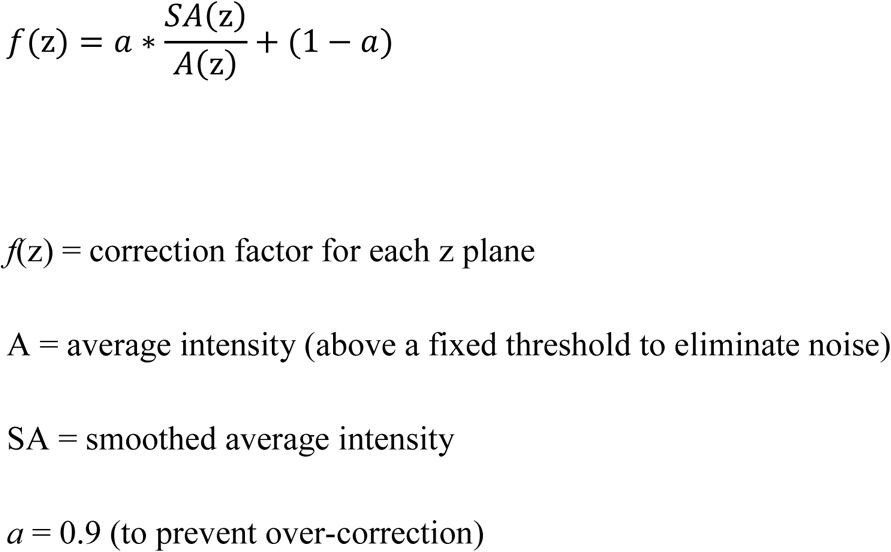

### Network extraction and quantification

#### Conduit network extraction

The voxel-based 3D EVIS image of LN conduits was processed to extract a connected conduit network suitable for 3D measurements. This was performed using a modified set of the tools we previously designed to isolate the blood vessel network from fluorescent 3D images of a mesenteric LN (33). In short, the grayscale 3D image is segmented using local thresholding, and the largest connected object selected to be skeletonized, followed by applying a tracing algorithm that transforms information from the segmented image and its skeleton into a topology map. This procedure generates a description of the network as a collection of connected tube segments, together with additional files allowing 3D visualization and manipulation. The image processing parameters were chosen specifically to allow for capturing fine conduit channels within the TCZ, while medullary regions with a high intensity of staining were excluded. To further narrow down the selection of TCZ conduits, B cell regions near the surface of the LN were manually removed using the filament editor in Amira (Thermo Fisher Scientific).

#### Exclusion of blood vessel surrounding conduit sleeves

One feature of the conduit network inevitably created a challenge for the processing: large blood vessels are often completely surrounded by conduits, resulting in the formation of conduit sleeves, hollow tubes which cannot be reduced to a single centerline by the skeletonization algorithm (S3 Fig). Previous studies have manually excluded these parts of the network (30), but given the large size of the present network we required a more automated approach in order to omit these sleeves. To this end we utilized the blood vessel image data from the same specimen, optimized using our previously described tools (33). We added them to the segmented image of the conduits, and filled remaining gaps manually and by using the segmentation tool in Amira, to obtain a combined 3D image of the conduits and blood vessels. As a result, conduits paths surrounding large blood vessels are reduced to the core blood vessel path helping to avoid artifacts and preserving the continuity of the network. Alternatively, the segmented image of the blood vasculature was subjected to ‘region growing’ in Amira and subtracted from the segmented image of the conduits, using a homemade tool that allows the addition and subtraction of pixel values between two images at the same location, in order to obtain a largely ‘blood vessel free’ conduit image. Both conduit datasets, either containing filled blood vessels or no blood vessels, were subjected to network extraction and topology analysis separately. Depending on the experimental question, we used either of these networks for the subsequent analysis as described accordingly (Fig 2 d-f).

#### Quantitative 3D measurements

The network topology map obtained from the extraction allows direct read-outs of network parameters, providing the number of segments, their volumes, the number of vertices per branch point, the branching angles, and length measurements. We used the blood vessel-free conduit topology map in this measurement, in order to obtain representative values for the conduit network without the contribution of blood vessels. In the calculation of branching angles, only segments with a length above 4 µm were chosen, to avoid the jittering artefact cause by very short segments. Additional tools were utilized to calculate the minimum distance from points in the network to the closest conduit (33), providing a measure of spacing of neighboring segments and allowing the gaps between them to be visualized as lit voxels. In this calculation all segments including potential blood vessels were assessed to avoid creating artificial gaps.

In order to investigate the possibility that there was a ‘preferred’ orientation of conduit segments in the network, a method was developed to estimate the tendency of conduits to align with a set of 13 directions roughly spanning the 3D range. The directions were chosen corresponding to the lines connecting a point in a regular 3D grid to its 26 nearest neighbors. The distribution of segment directions over these 13 reference directions was calculated by summing the magnitude of segment projections onto the 13 lines, then normalizing. The results were visualized in LabVIEW (National Instruments).

A ‘moving average’ display providing insight into the relative segment density was obtained by first computing the averaged voxel densities of cubes with a set radius (e.g. 10 μm) while moving in 2 μm steps across the binary volume image, then rescaling the density values from 0-1 to 0-255, and finally visualizing the resulting averaged 3D image as a greyscale or false-colored (e.g. heatmap) image using ImageJ (2D) and Amira (3D).

#### Selection of subregions

To specifically measure and compare anatomical differences between the outer and inner TCZs, spherical subregions with a diameter of 100 µm were selected for individual analysis from both zones. Since the identification of non-touching subregions within an irregular shaped 3D volume is not trivial and automated tools are lacking, we manually selected the center points for each of the subregions based on the observable segment density in z planes using ImageJ (NIH). By specifying a center point and radius in a 3D cropping tool, these regions of interest could be isolated and their topology determined individually. As above, the topology map exclusive of blood vessels was used to obtain conduit-specific parameters but blood vessels were included to estimate the distance distribution to the nearest conduit.

#### Modelling T cell motility

Based on the current understanding that the FRC network provides a substrate for T cell migration, we sought to simulate T cell motility on the 3D conduit network. The coefficient of motility, *C*_*m*_, which is analogous to a diffusion coefficient (74), was estimated by simulating the movement of a large number of cells on the network, subject to certain assumptions about speed and behavior at junctions. The procedure is as follows. A large number of cell paths through the network are simulated, the starting point (and starting direction) of each path chosen at random. Each cell is initially assigned a speed drawn from a Gaussian distribution with specified mean (here: 13 µm min^-1^) and coefficient of variation - (standard deviation)/mean (here: 0.1). The cell moves with this constant speed along the network segments. When a segment junction is encountered the branch taken by the cell is determined randomly, according to the following procedure. For each possible branch, *k*, the turning angle *θ* is determined, and for an angle less than 90° the probability weight *w*(*k*) associated with that branch is computed as cos^4^(*θ*), the fourth power of the cosine of the turning angle, otherwise *w*(*k*) is set to a very small value (0.001). The probability of taking branch *k* is then given by *w*(*k*) divided by the sum of all the weights. The actual branch taken is then determined in the usual way by generating a random variate with a uniform distribution. If a cell reaches a dead-end in the network the direction of movement along the segment is reversed. To reduce the encounter of dead-ends which could skew the observed *C*_*m*_, each tested network initially underwent a healing step of pruning and joining dead-ending segments to neighboring vertices with a maximum branch length of 15 µm.

In short, if the junction-directed unit vector corresponding to the branch that the cell is on is *v*(0), and there are *N*_*b*_ branches the cell can take, with unit vectors [*v*(*k*),*k*=1,..,*N*_*b*_] directed away from the junction, then the turning angle onto the *k*th branch is given by the dot-product of two unit vectors (cos^-1^ is the inverse cosine function.):

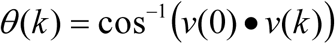

Then the probability of taking branch *k* is given by:

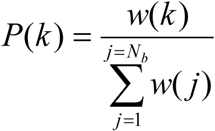

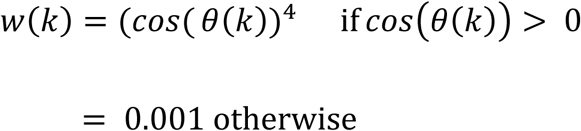

The movement of each of 5000 cells across the network was simulated in this way for a period of one hour. In order to avoid the possibility of a cell reaching the boundary of the network the starting points were restricted to those falling within a sphere of radius 100 μm centered at the center of the LN, unless otherwise specified for TCZ subregions. The average squared distance of cells from their start points was computed at 5 min intervals, and plotted. The estimate of the coefficient of motility (*C*_*m*_) is the slope of the resulting line divided by 6. Since the curve is not a straight line the average slope was estimated from the points at times 0 and 60 min (= T).

Let (*x*_i_ (*t*), *y*_i_ (*t*),*z*_i_ (*t*)) be the position of cell *i* at time *t, N* = number of cell paths simulated.

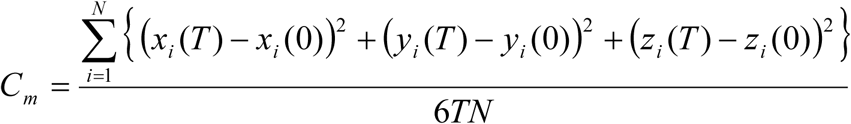

To account for the tissue shrinkage that occurs during the embedding process (estimated to be 20% in each direction), a correction factor of 1.25 can be applied to all segment lengths in this simulation.

A selection of 20 tracks, normalized to the same starting point, was visualized in a spider plot using CMGUI. Calculation of *C*_*m*_ was performed on the conduit network in which blood vessels were removed and on the full network including blood vessels, to account for the possibility that blood vessels can serve as additional migration paths.

#### Visualization

3D images obtained from extended-volume or conventional confocal imaging were acquired as greyscale 3D tiff files in raw format and were pseudo-colored, processed, and superimposed with the 3D rendering programs Voxx, ImageJ (NIH), Amira (Thermo Fisher Scientific), or Imaris (Bitplane). Visualization of 3D image data was performed by generating 2D projections of rendered volume images, isolating and displaying single z-planes, or by accumulating several z planes over a range of 10-20 µm to provide ‘thick volume sections’ that can allow better insight into the arrangement of fine structures over a restricted range of tissue. Selected programs such as Voxx and Imaris further allowed the generation of high quality movie files. The 3D rendering software CMGUI was employed for rendering the network and the simulated cell paths. Graphs were generated in GraphPad Prism version 7.02 for Windows (GraphPad Software).

### Quantification and Statistical Analysis

All statistical analysis was performed in GraphPad Prism v7.03. Statistical parameters including the exact value of N, the definition of center, dispersion and precision measures (mean ± SD) and statistical significance are reported in the Fig 3 and the respective figure legend. Data is judged to be statistically significant when p < 0.05 by two-tailed Student’s t test. In figures, asterisks denote statistical significance as calculated by Student’s t test (*, p < 0.05; **, p < 0.01; ***, p < 0.001; ****, p < 0.0001; ns = not significant).

### Data and Code Availability

A custom-written toolset to extract volumetric network information from a voxel-based 3D image was modified from our previous work (33) to allow for the isolation of a connected conduit network in the T cell zone. The source code for these tools can be found under: https://github.com/gibbogle/vessel-tools.git. The 3D data presented in this study are available from the corresponding author upon reasonable request.

## Supporting information

Supplemental Images S1-S4

## Acknowledgments

The authors gratefully acknowledge the help and advice of Dr Adrian Turner, Mrs Amorita Petzer, Ms Shorena Nachkebia, and the members of the Dunbar and LeGrice laboratories at the University of Auckland. We thank the Biomedical Imaging Research Unit (BIRU) at the University of Auckland for providing access to the image processing software Amira. G.B. acknowledges the support of the Auckland Bioengineering Institute. This work was partially funded by the Maurice Wilkins Centre, a national Centre of Research Excellence hosted by the University of Auckland.

## Author Contributions

P.R.D., I.D.K. and G.B. conceived and designed the study. I.D.K. conducted the experiments and analyzed the data. A.P. supervised the surgical experiments. G.B.S. supervised the EVIS imaging and image processing, and developed targeted tools to assist the image processing. G.B. developed computational tools and supervised the analysis. I.D.K., G.B. and P.R.D. wrote the manuscript. G.B.S., A.P. and I.J.L. gave technical support and conceptual advice on the project.

## Supplementary Items Titles

S1-S5 Fig.

S1 Video. Fly-through animation of an entire murine LN captured by EVIS imaging. 3D image reconstruction of this dataset visualizes the lymphatic (red glow) and blood (green) passageways in a slice-by-slice view moving through z sections of 20 µm thickness and provides an interior view of LN sub-compartments including the staining-rich medulla, a dense mesh of conduit channels in the central TCZ, and the B cell follicles emerging near the SCS at the rim of the LN. Image reconstruction and animation was performed in Voxx. Related to Fig 1.

S2 Video. This fly-through animation is moving slice-by-slice though the 3D volume image of a murine LN in optical z sections of 20 µm thickness, zoomed into the interface between the paracortex and the medulla. Blood vessels (green) penetrate through the LN volume rich in lymphatic staining (red glow), each surrounded by a conduit sleeve. Conduit channels frequently terminate on cortical sinuses continuous with the lymphatic system (sinuses) of the medulla. The dataset was acquired using EVIS imaging and visualized in Voxx. Related to Fig 1.

S2 Video. 3D surface representation and animation of the LN blood vessels (red), lymphatic sinuses (green), and conduit channels (grey). A crop from the edge of the paracortical region of a WGA-perfused murine LN exemplifies the tight relationship of the blood and lymphatic passageways within the LN as conduit channels meet a plexus of lymphatic sinuses. Several blood vessels can be seen enclosed by a sleeve of conduits. The 3D image data were generated using EVIS imaging, lymphatic sinuses were isolated from the conduit data with the help of custom image processing tools, and surface rendering and animation of the data was performed in Imaris. Related to Fig 2.

S4 Video. 3D reconstruction and animation of the blood vessel system (red) and lymphatic channels (white) of a murine LN. In the first part of the animation the blood vasculature is shown in full and rotated around the Y axis followed by a slice-by-slice view of the LN moving through z sections of 10 µm thickness and displaying both the lymphatic and blood vessel anatomy. Here, accumulations of the fluorescent tracer (WGA) used to visualize the lymphatic passageways (white) can be observed within the B cell follicles, which appear as spherical structures below the SCS devoid of an organized conduit network. Within these locations the tracer material is labelling interconnected clusters resembling the FDC network. The 3D image was acquired using EVIS imaging and reconstructed and animated in Imaris. Related to Fig 4 a.

S5 Video. A close-up view of a ‘follicular reservoir’ (white) within a B cell follicle labelled with WGA within a murine LN. A 3D reconstructed image of the conduit system near the SCS is reduced slice-by-slice to open the view to a spherical cluster with strong fluorescent label, followed by building the image up again in a slice-by-slice manner. Standard confocal microscopy was performed with a voxel resolution of 0.36 × 0.36 × 1 µm over a depth of 40 µm followed by image reconstruction and animation in Voxx. Related to Fig 4 g, h.

