## Supplemental Images S1-S4 for "High-resolution 3D imaging and topological mapping of the lymph node conduit system"

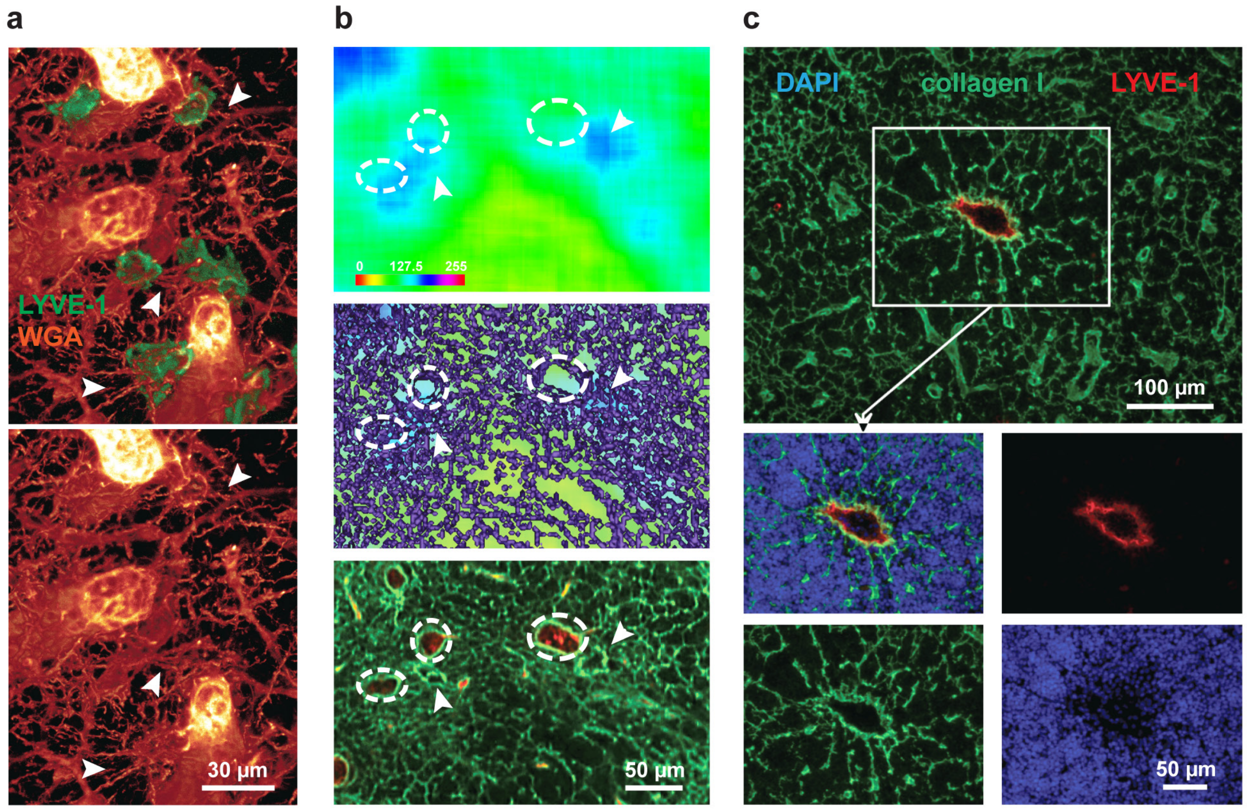


**S1 Fig. Conduits channels focus on lymphatic sinuses. Related to Fig 1.**

High-resolution confocal images of LN conduits, blood vessels (both co-labelled via perfusion in red) and lymphatic sinuses (LYVE-1, green) display conduits fusing into sinuses in close proximity to large blood vessels (arrowheads, **a**). In a moving average display (upper panel) generated from 3D LN images with WGA-labelled lymphatic channels that color-codes pixel density in a rainbow spectrum, regions with a high density of conduits appear blue (**b**). These blue regions do not overlap with the location of blood vessels, seen as gaps in the conduit mesh (dashed circles), as can be taken from an overlay of the moving average image with the corresponding section of the conduit network image (middle panel, **b**). Instead, regions with high density of conduit stain show an association with lymphatic sinuses (arrowheads) that stain brightly with WGA (green) and lack a vascular core (red, lower panel, **b**). In multicolor fluorescent images of an immuno-labelled LN section, several collagen I+ conduits concentrate on a LYVE-1+ lymphatic sinus (**c**).

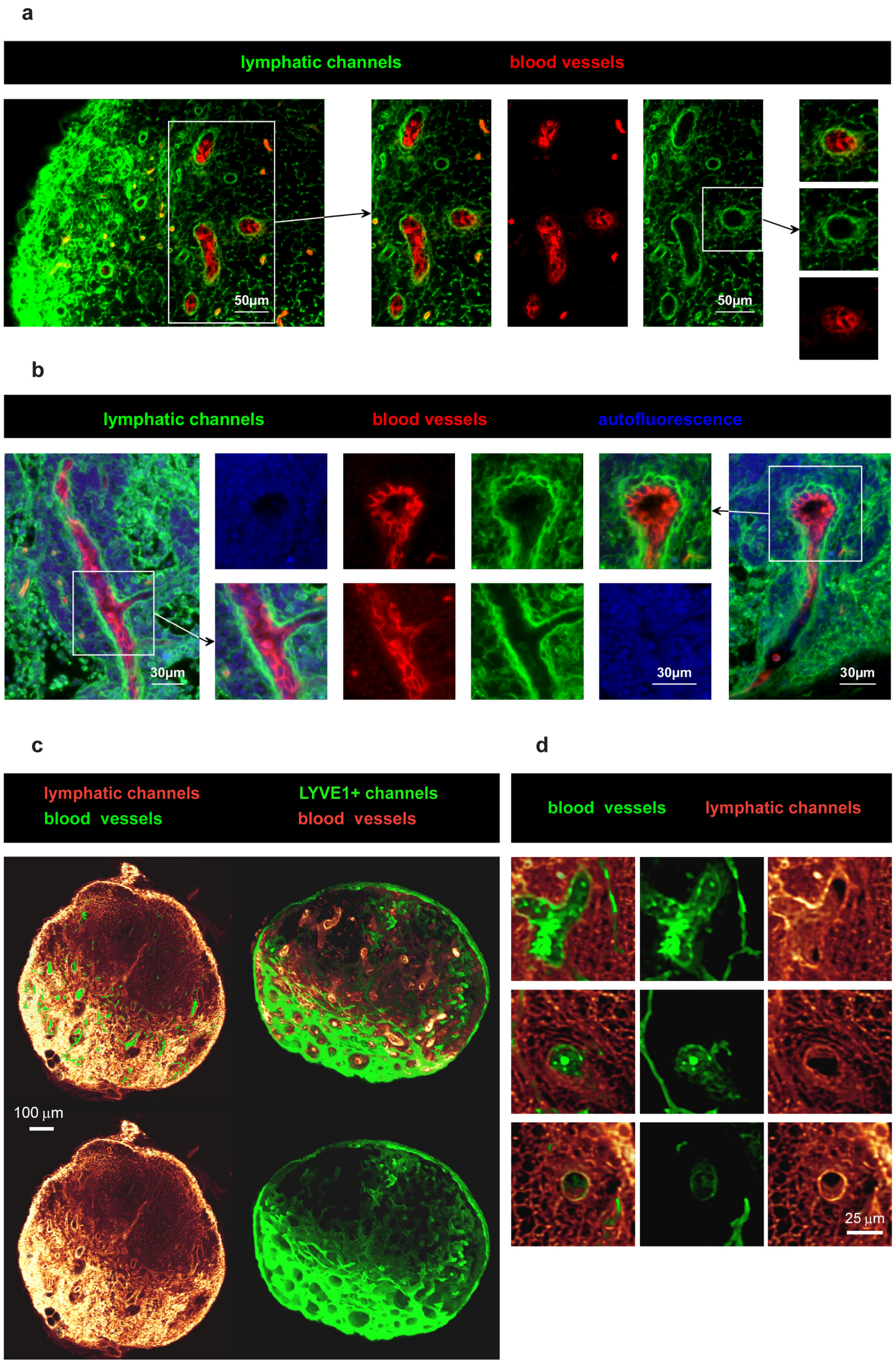

a
lymphatic channels
blood vessels
50µm
50µm
b
lymphatic channels
blood vessels
autofluorescence
30µm
30µm
30µm
c
lymphatic channels
blood vessels
LYVE1+ channels
blood vessels
100 µm
d
blood vessels
lymphatic channels
25 µm

**S2 Fig. The conduit network forms sleeves around blood vessels. Related to Fig 2.**

Murine LNs were perfused with fluorescently tagged WGA and 2000 kDa dextran to label the blood vasculature (red) and locally injected with fluorescently tagged WGA to label the lymphatic channels including conduit passageways (green). Multicolor fluorescent images of 2 µm LN sections show LN blood vessels are surrounded by sleeves continuous with the conduits (**a**). Close-up fluorescent images confirm the close juxtaposition of blood vessel endothelium (red) enclosed by a cell layer stained with lymph-borne WGA (green) against the background of autofluorescent cell bodies (blue, **b**). 3D reconstructed images of a LN volume image generated by EVIS imaging (at 1 µm pixel resolution, **c**) visualise the overall arrangement of the WGA-labelled channels including conduits and lymphatic vessels (red glow, left panel), the latter of which also stain positively for LYVE-1+ (green, right panel), against the dense network of blood vessels weaving through the LN. Close-up images of 20 µm optical sections of a LN volume image illustrate how the conduit sleeves (red glow) fully enclose blood vessels (green, **c**).

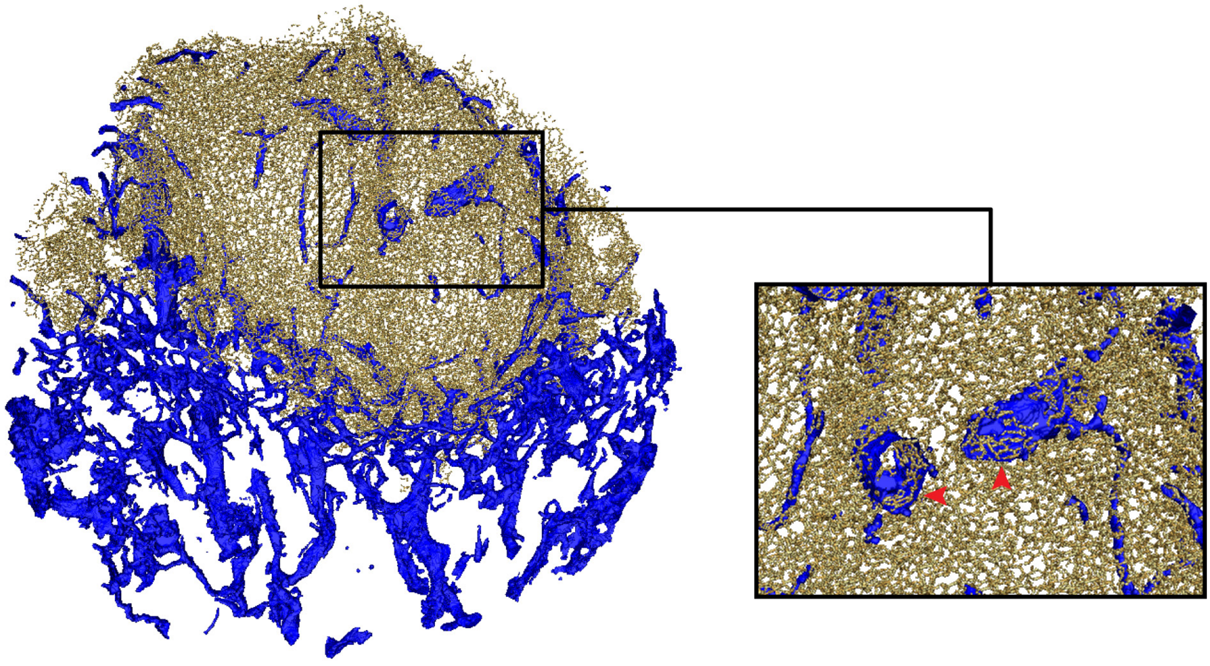

**S3 Fig. Conduit network extraction produces unavoidable artefacts. Related to Fig 2.**

A volume projection of the LN blood vasculature (blue) and conduit network (gold) demonstrates typical artefacts that occur during the segmentation of the fine conduit network around large blood vessels (close-up box). Here, the conduit network encloses the blood vasculature entirely and forms large tubes or sleeves that cannot be interpreted by the skeletonisation algorithm, resulting in the creation of many short segments along the conduit sleeve (red arrowheads), hindering realistic analysis of the network at these locations.

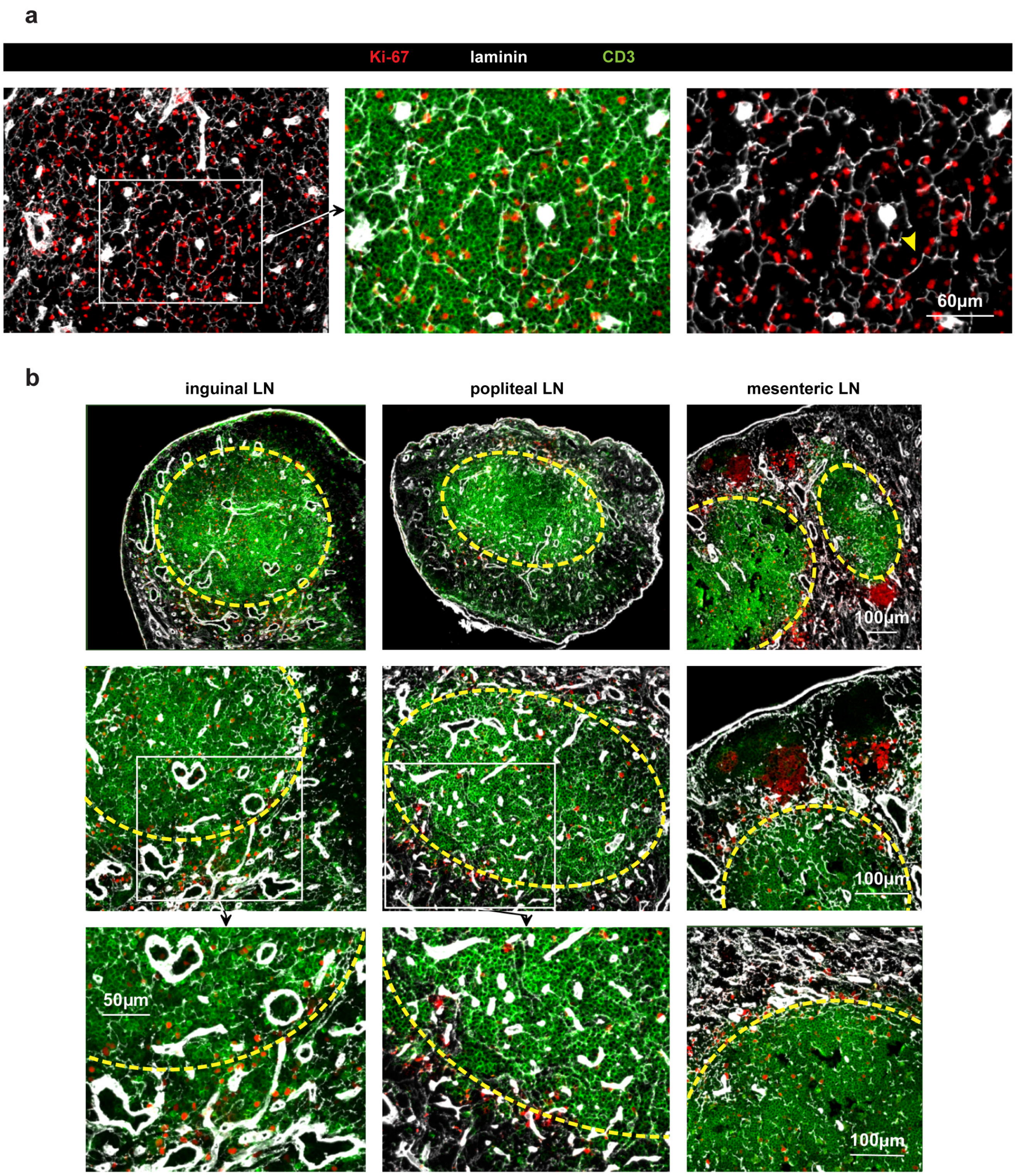


**S4 Fig. Proliferating T cells in LNs are located close to conduits and accumulate in the superficial TCZ. Related to Fig 3.**

Multicolor fluorescent images of immuno-labelled LN sections reveal a close association between Ki-67+ cells and laminin+ conduits in the CD3+ TCZ of an inguinal LN (yellow arrowhead) (**a**). Overview of CD3+ TCZs (green) in inguinal, popliteal, and mesenteric LNs in which Ki-67+ cells can often be found close to the border of the TCZ (yellow dashed line, **b**).
